# Layered double hydroxide nanoparticles induce composition-dependent cytotoxic and phenotypic effects in mammalian cells

**DOI:** 10.64898/2026.07.28.741015

**Authors:** André Lopes Ferreira, Luana Pereira Cardoso, Thaís Moraes-Lacerda, Leticia Ester dos Santos, Lillia Iamar Leite Maciel Gama, William Reis de Araujo, Marcelo Bispo de Jesus

## Abstract

Layered double hydroxides (LDHs) are increasingly explored for agricultural, environmental, and biodelivery applications, but their composition-dependent effects on mammalian cells remain insufficiently defined. Here, we synthesized Al–Ni, Al–Co, and Al–Cu LDH nanoparticles and evaluated their physicochemical properties and biological responses across exposure-relevant mammalian cell models. The formulations showed hydrodynamic diameters of approximately 200–300 nm, moderate dispersity, strongly positive surface charge, and characteristic lamellar LDH features. Cytotoxicity was assessed using MTT, Calcein-AM, and Hoechst–PI assays in HaCaT, A549, and HT-29 cells, representing dermal, pulmonary, and intestinal exposure contexts, together with NIH/3T3 fibroblasts as a sensitive comparative model. LDH toxicity was strongly dependent on metal composition and cell type, with an overall trend of Al–Cu > Al–Co > Al–Ni and more pronounced cytotoxic effects in A549 and HT-29 cells. To detect cellular perturbations beyond overt viability loss, we applied high-content imaging using Live Cell Painting. Multiparametric single-cell profiling revealed composition- and dose-dependent alterations in acidic vesicle organization, nuclear texture, and cytoplasmic granularity. Notably, phenotypic deviations were detected at concentrations below those producing measurable effects in conventional viability assays, and linear discriminant analysis separated the phenotypic signatures induced by the three LDH formulations. Together, these findings show that LDH biological activity cannot be generalized across metal compositions and that high-content phenotypic profiling provides added sensitivity for detecting early cellular perturbations. This integrated approach supports composition-aware nanosafety evaluation and may inform the safer development of LDH-based technologies for agricultural and biotechnological applications.

## 1. Introduction

The global agricultural sector faces increasing pressure to expand productivity while reducing environmental and health impacts associated with intensive chemical inputs. Projections indicate that the world population will reach approximately 9.2 billion people by 2050, driving an estimated 98% increase in food demand^1^. To support this trend, agricultural productivity must rise substantially to balance supply and demand. In most production systems, this growth has been supported by the intensive use of chemical inputs, including fertilizers, herbicides, and pesticides, which enhance crop yields and generate immediate economic benefits. However, their widespread and often indiscriminate use has resulted in the accumulation of toxic residues in soil, water, and air^2,3^. These environmental burdens, combined with high financial costs related to fertilization, irrigation, and energy consumption, as well as adverse effects on human health and degradation of arable land, threaten the long-term sustainability of global agriculture^1,4^. Collectively, these pressures call for strategies that increase agricultural productivity and product quality while minimizing environmental impacts and input costs.

Among the emerging alternatives, layered double hydroxides (LDHs) have gained increasing attention. LDHs are nanostructured materials composed of divalent and trivalent metal cations arranged in a lamellar structure, with interlayer spaces containing anions (*e.g.*, nitrate, sulfate, carbonate) and water molecules^5,6^. Their general formula, [M^II^_1-x_M^III^_x_(OH) _2_] *^x^* ^+^ (A *^n^* ^−^) *_x_* _/_ *_n_* · *m* H _2_ O, allows broad chemical tunability through variations in metal composition and interlayer anions^7,8^. Owing to their structural versatility and physicochemical properties, LDHs have been explored as catalyst supports^9,10^, nanocomposites^11,12^, gas adsorption^13^, drug delivery^14–16^, antibacterial applications^17,18^, nutrient storage, and controlled nutrient release^7,8^. In agriculture, LDHs have been investigated for nutrient storage and delivery^19–22^, environmental remediation^23^, and as a platform for gene editing and delivery^24,25^. Their versatility, stability, and scalability therefore position LDHs as promising materials for emerging agricultural and biotechnological technologies.

Despite their widespread use and potential for agricultural innovation, there is a notable lack of systematic studies addressing the toxicity of LDH nanoparticles across the major human exposure routes: inhalation, dermal penetration, and ingestion. This gap is concerning because LDHs intended for agricultural applications can reach the environment and the human body through multiple exposure routes. Nanoparticles dispersed in air may reach the respiratory tract, where ultrafine particles (<2.5 μm) have been epidemiologically associated with increased pulmonary morbidity and mortality^26,27^. Similarly, dermal exposure may occur through occupational handling or contaminated surfaces, allowing nanoparticles to penetrate the stratum corneum via intercellular, intracellular, or transappendageal routes^28–30^. Ingestion is yet another relevant pathway, as nanoparticles can reach the gastrointestinal tract through contaminated food, water, or swallowed airborne particles, interacting with intestinal epithelial cells and potentially affecting barrier integrity^31^. In particular, little is known about how variations in divalent metal composition affect cytotoxicity and sublethal cellular perturbations across biologically relevant mammalian models^32^. This knowledge gap complicates the safe development and risk assessment of LDH-based technologies.

To address this knowledge gap, we synthesized and comparatively evaluated Al–Ni, Al–Co, and Al–Cu layered double hydroxide (LDH) nanoparticles using an integrated physicochemical and biological approach. The formulations were characterized with respect to their structural, morphological, and colloidal properties prior to toxicological assessment. Biological responses were evaluated in mammalian cell lines representing exposure-relevant tissues, including HaCaT keratinocytes, A549 alveolar epithelial cells, HT-29 intestinal epithelial cells, and NIH/3T3 fibroblasts as a comparative murine model^33,34^. Cytotoxicity was assessed using orthogonal viability assays, while high-content imaging and immunocytochemical analyses were applied to capture sublethal phenotypic and stress-associated cellular alterations. By integrating conventional toxicology with multiparametric phenotypic profiling, this study investigates how LDH metal composition influences mammalian cellular responses and provides experimental support for composition-aware nanosafety assessment of LDH-based technologies.

## 2. Materials and Methods

### 2.1 Synthesis of Layered Double Hydroxide Nanoparticles (LDHs)

For LDH synthesis, 10 mL saline solutions were prepared using divalent metal cations (Ni^2+^, Co^2+^ and Cu^2+^, 0.70 mol/L) and trivalent metal cations (Al^3+^, 0.30 mol/L) in the desired proportions. This mixed solution was slowly added to 40 mL of reaerated NaOH (0.45 mol/L) under a nitrogen atmosphere and vigorous stirring for 10 min. The resulting LDH paste was collected by centrifugation (6,500 RPM), washed 2 times, and manually redispersed using a vortex in 80 mL of deionized water. The dispersion was then hydrothermally treated in an autoclave at 100 °C for 16 h, yielding a homogeneous and stable LDH suspension.

### 2.2 Physicochemical Characterization

#### 2.2.1 Morphological Characterization

Layered double hydroxide (LDH) nanoparticles were characterized by transmission electron microscopy (TEM) to evaluate their morphology, size, and structural organization. For the analysis, the samples (0.05 mg/mL) were deposited onto copper grids coated with a formvar film and incubated with a 1% (w/v) uranyl acetate solution to provide contrast. Images were acquired using a Tecnai G2 Spirit BioTWIN transmission electron microscope operated at an accelerating voltage of 80 kV.

#### 2.2.2 Dynamic Light Scattering (DLS)

Dynamic light scattering (DLS) was used to determine the hydrodynamic diameter, polydispersity index, and zeta potential of the nanoparticles using a ZetaSizer Nano ZS 90 (Malvern, UK). Measurements were performed at 25 °C in polystyrene cuvettes with a 10-mm path length at a scattering angle of 90° and presented as a percentage of intensity. Zeta potential was measured using capillary cells with the same path length. All LDH suspensions were standardized at 0.1 mg/mL in water; analyses were performed 3 times on independent samples.

#### 2.2.3 Fourier Transform Infrared Spectroscopy (FT-IR)

Structural characterization was performed using FT-IR spectroscopy (Nicolet Summit IR 200 FT-IR) in reflectance mode with 120 scans, 4.0 cm□¹ nominal resolution, and a spectral range of 500–4000 cm^-1^.

#### 2.2.4 X-ray Diffraction (XRD)

X-ray diffraction patterns were obtained using a Shimadzu XRD-7000 diffractometer equipped with a Ni filter and Cu Kα radiation, with a scanning rate of 2°/min and a current of 50 mA.

### 2.3 In vitro biological assessment of LDHs

#### 2.3.1 Cell Culture

To evaluate the cytotoxicity of LDH nanoparticles across biologically relevant exposure routes, we employed three mammalian cell lines: human keratinocytes (HaCaT), human colon adenocarcinoma cells (HT-29), and human alveolar basal epithelial cells (A549) together with a comparative mouse embryonic fibroblasts (NIH/3T3) model. These lines represent dermal, pulmonary, and gastrointestinal exposure pathways and are widely used in nanosafety research, providing biologically relevant models for early hazard identification. All cell lines were cultured in Dulbecco’s Modified Eagle Medium (DMEM; Gibco) supplemented with 10% fetal bovine serum (FBS; Gibco, South America) and 1% Penicillin–Streptomycin (Gibco, Germany). Cells were maintained in a Panasonic humidified incubator at 37 °C, 95% relative humidity, and 5% CO_2_. Routine subculturing was performed at 70–80% confluence using 0.25% trypsin-EDTA (Thermo Fisher Scientific). Cells were never maintained in continuous culture for more than two months, and all experiments were conducted between passages 5–20, depending on the cell line.

To ensure culture integrity, cells were routinely screened for mycoplasma contamination every two weeks and prior to experimentation. Testing was performed on antibiotic-free cultures for at least 72 h using Hoechst 33342 DNA staining followed by fluorescence microscopy. Only mycoplasma-negative batches were used for experiments.

#### 2.3.2 Cell Seeding and Exposure Conditions

For cytotoxicity experiments, each cell line was seeded at a density optimized for its growth rate and morphology. Cells were seeded at 5.0×10^3^ (NIH/3T3), 4×10^3^ (HaCaT), 2×10^2^ (HT-29), and 2×10^3^ (A549) per well in 96-well plates. After seeding, plates were incubated for 24 hours at 37 °C, 5% CO_2_, and 95% relative humidity to allow cell attachment. Background controls (medium only and untreated cells) were included in all plates to ensure proper baseline correction for absorbance- and fluorescence-based assays. Following the 24-h recovery period, the culture medium was replaced with fresh medium containing nominal LDH concentrations ranging from 1.95 to 500 µg/mL. Cells were exposed for an additional 24 hours under standard culture conditions. Unless otherwise specified, all cytotoxicity assays were performed using three technical replicate wells per condition in three independent biological experiments.

#### 2.3.3 MTT Assay

Following 24 h of LDH exposure, the medium was replaced with 1.0 mg/mL MTT solution prepared in serum-free DMEM; untreated cells were used as a control. Plates were incubated for 3 h at 37 °C to allow formazan crystal formation. The MTT solution was replaced with 100 μL DMSO to each well to solubilize the formazan. Plates were gently agitated for 5 minutes, and absorbance was measured at 570 nm using a Cytation 5 Hybrid Multi-Detection Reader (BioTek Instruments, USA).

#### 2.3.4 Calcein-AM Assay

Following 24 h of LDH exposure, the medium was replaced with FluoroBrite™ DMEM containing Calcein-AM (0.025 µmol/L) for 30 min at 37 °C; untreated cells were used as a control. Fluorescence was measured (excitation 494 nm, emission 517 nm) using a Cytation 5 Hybrid Multi-Detection Reader (BioTek Instruments, USA).

#### 2.3.5 Imaging-based Viability Assay: Calcein-PI-Hoechst

Following 24 h of LDH exposure, the medium was replaced with a staining solution containing Calcein-AM (0.025 µmol/L), propidium iodide (PI, 1 µg/mL), and Hoechst (1 µg/mL). After 30 min incubation, plates were imaged in six fields per well using a 10× objective on the Cytation 5 system, with the following filter sets: GFP (EX 469/35 nm, EM 525/39 nm, dichroic mirror 497 nm, LED 465 nm) for Calcein-AM; PI (EX 531/40 nm, EM 647/57 nm, dichroic mirror 605 nm, LED 523 nm) for PI; DAPI (EX 377/50 nm, EM 447/60 nm, dichroic mirror 409 nm, LED 365 nm) for Hoechst 33342. Hoechst-positive nuclei represented total cells, while PI-positive nuclei represented dead cells. Image acquisition was performed using identical exposure settings across all conditions to ensure comparability. Cell segmentation and counting were performed using the Gen5 3.12 software (BioTek), with Hoechst used to quantify total nuclei and PI to identify dead cells. Calcein-AM fluorescence was used for qualitative visualization of viable cells. Cell viability was calculated by subtracting the number of PI-positive nuclei from the total Hoechst-positive nuclei, and results were expressed in two ways as a percentage relative to the control group using the following equation:

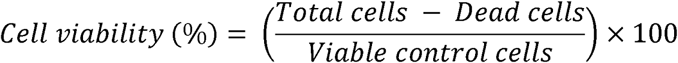

#### 2.3.6 High-Content Image Analysis: Live Cell Painting Assay

To assess cellular alterations induced by LDH nanoparticles beyond those detected by conventional viability assays, high-content imaging was performed using the Live Cell Painting assay^35,36^. Nominal exposure concentrations were selected based on the IC_50_ values determined in MTT, and included IC_50_, IC_50_/10, and IC_50_/100 for each LDH formulation. Antibiotic- and serum-free DMEM was used as the negative control, lactose (2 µM, 100 µL; Sigma-Aldrich, USA) as an additional reference condition. Phenotypic profiles were generated from 3 technical replicate wells per condition across 3 independent biological experiments; untreated cells were used as a control.

After 24 h of treatment, the medium was removed and replaced with 100 μL of Acridine Orange (AO, 10 μM) prepared in FluoroBrite™ DMEM. After 10 minutes of incubation at 37 °C, the staining solution was replaced with 100 μL FluoroBrite™ DMEM and fluorescence images were acquired at 12 sites per well using a Cytation 5 equipped with a 20× Olympus Plan Fluorite objective (NA 0.45). The following filter cubes were used: GFP (EX 469/35 nm, EM 525/39 nm) for AO–nucleic acid signal; PI (EX 531/40 nm, EM 647/57 nm) for AO–acidic vesicle signal; DAPI (EX 377/50 nm, EM 447/60 nm) for Hoechst (when nuclear labeling was required for segmentation in specific cell lines, 3T3, A549 and HT-29).

#### 2.3.7 Image segmentation and feature extraction

Image quality was assessed using CellProfiler by evaluating blur and intensity metrics. Images failing to meet image quality thresholds (established in CellProfiler Analyst) were excluded from further analysis. A CellProfiler pipeline was then developed to segment multiple cellular compartments, including nuclei, whole cells, cytoplasm, nucleoli, and vesicles. Nuclear segmentation was performed using the RunCellpose plugin, which applies pre-built Cellpose models^37^. Before full-scale analysis, the pipeline was tested on one representative image per well from each plate to detect and correct potential segmentation issues. Segmentation accuracy was verified by overlay inspection^37,38^. Following segmentation, morphological, intensity, and texture features were extracted for each object using CellProfiler modules^39^. These features included morphological descriptors (*e.g.*, area, perimeter, roundness), intensity features (*e.g.*, integrated and mean pixel intensity across channels and compartments such as the cell, nucleus, and cytoplasm), and texture features capturing object- and image-level roughness and homogeneity^40^. Radial distribution analysis was also performed, dividing each cell into four concentric bins to quantify GFP-AO and PI-AO intensity gradients. All per-cell outputs were consolidated using pycytominer, ensuring a standardized and reproducible data structure for downstream profiling^41^.

#### 2.3.8 Profile generation, normalization, and multivariate analysis

To derive meaningful profiles from the extracted features^42^, we processed image-based data using Pycytominer^41^. Single-cell measurements were aggregated into well-level profiles using the median function within the aggregate module. Metadata annotations included compound identity, nominal concentration, and experimental batch. Feature normalization was performed on a per-plate basis relative to negative controls using the RobustMAD transformation (scaled feature = (x – median)/MAD). Features were subsequently filtered to remove variables with more than 5% missing values, low variance, pairwise correlations exceeding 0.9, or extreme outliers (> 500). The resulting curated dataset was used for all downstream analyses. Dimensionality reduction was performed using Linear Discriminant Analysis (LDA) on the normalized well-level profiles to visualize treatment-dependent phenotypic separation. Unsupervised clustering was performed using the k-means algorithm to identify groups of treatments with similar phenotypic signatures.

#### 2.3.9 Immunofluorescence Assays

Cells were seeded at standard densities in black 96-well plates and exposed for 24 h to nominal LDH concentrations corresponding to the IC_50_ of each cell line, as well as 50, 150, and 250 µg/mL. After incubation, cells were fixed with 4% paraformaldehyde, washed with PBS, permeabilized with 0.3% Triton X-100, and blocked with 5% goat serum for 1 h. Cells were incubated with primary antibodies against Ki-67 (8D5) and γ-H2AX (10hosphor-Ser139) (Merck, Germany) for 1 h at 37 °C. After washing, cells were incubated with Alexa Fluor™ 546 goat anti-mouse, Alexa Fluor™ 488 goat anti-rabbit, Hoechst 33342, and Alexa Fluor™ 647 Phalloidin for 1 h at room temperature in the dark. Imaging was performed on a Cytation 5 using GFP, RFP, DAPI, and CY5 filters at 20× magnification. Image segmentation was performed in CellProfiler, with nuclear and cytoplasmic segmentation assisted by CellPose using Hoechst and Phalloidin channels.

### 2.3 Statistical analysis

For cytotoxicity assays (MTT, Calcein-AM, and Calcein-PI-Hoechst), cell viability and cell count values were normalized to untreated controls to calculate the percentage of viable cells. The half-maximal inhibitory concentration (IC_50_) values were determined using nonlinear regression analysis in GraphPad Prism (version 8.0.2). For immunofluorescence analysis, comparisons of protein fluorescence intensity were performed between nanoparticles within each concentration. Statistical analysis was conducted using two-way ANOVA followed by Tukey’s post hoc test for multiple comparisons. All experiments were conducted in biological triplicate (n = 3). Differences were considered statistically significant when p < 0.05.

## 3. Results

### 3.1 Al–Ni, Al–Co, and Al–Cu LDHs display controlled physicochemical properties and characteristic lamellar structure

A broader set of Al-based LDH formulations was initially synthesized and screened for colloidal properties. From this screening, three formulations (Al–Ni, Al–Co, Al–Cu) were selected for further analysis based on their colloidal stability and suitability for biological evaluation **(Table 1)**. Dynamic light scattering revealed that Al–Ni, Al–Co, and Al–Cu formulations exhibited hydrodynamic diameters ranging from approximately 184 to 327 nm with moderately uniform dispersity (PDI 0.19–0.33) (Table 1). All three selected particles displayed strongly positive zeta potentials, ranging from +33 to +46 mV **(Table 1)**. FTIR analysis confirmed that each composition preserved the hallmark features of LDH materials, including the broad O–H stretching band (∼3440–3450 cm□¹), the H–O–H bending band (∼1620 cm□¹), the interlayer anion band (∼1380–1390 cm□¹), and metal–oxygen and metal–hydroxide stretching modes between 580–870 cm□¹ **(Figure 1A–C)**^43^. These spectral signatures were maintained across Ni^2+^-, Co^2+^-, and Cu^2+^-containing LDHs. X-ray diffraction patterns further confirmed the lamellar structure of the LDHs, showing the characteristic basal reflections (003), (006), and (009)^44^ **(Figure 1G–I)**. In addition, transmission electron microscopy images revealed platelet-like particles with well-defined hexagonal morphology **(Figure 1J–L)**.

**Table 1.** Physicochemical characterization of aluminum–based LDH formulations containing different divalent ions (n=5).

| Composition<br>(M<sup>2+</sup> / M<sup>3+</sup>) | Size (nm) | PDI | Result<br>quality | Zeta potential<br>(mV) |
| --- | --- | --- | --- | --- |
| Al–Ni | 184.6 ± 23.74 | 0.218 | Good | +39.2 ± 1.79 |
| Al–Co | 327.3 ± 12.8 | 0.193 | Good | +33.4 ± 1.18 |
| Al–Cu | 273.3 ± 25.56 | 0.333 | Good | +44.2 ± 2.58 |

**Figure 1.**
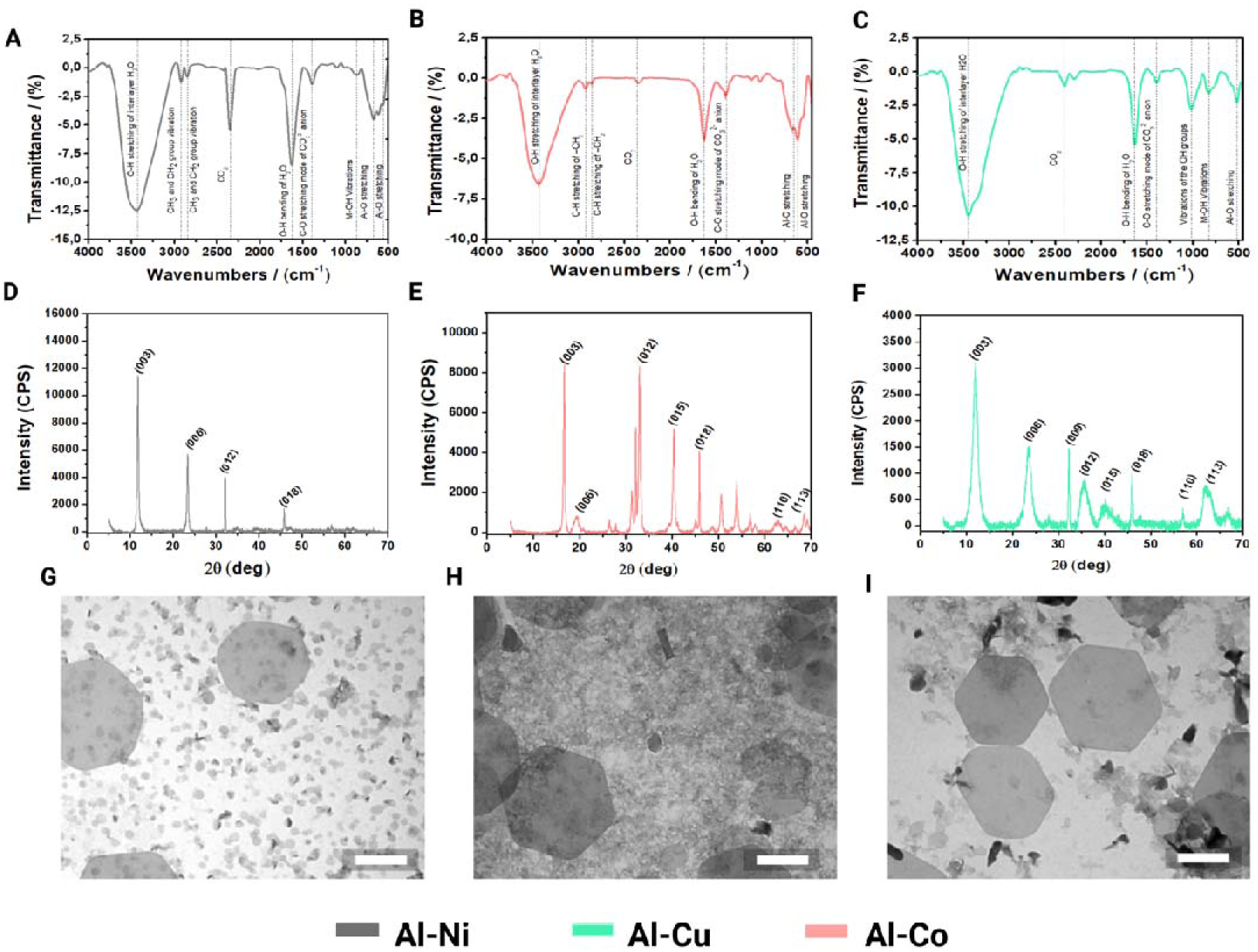
Structural and physicochemical characterization of Al–Ni, Al–Co, and Al–Cu LDHs. (A–C) FTIR spectra of Al–Ni, Al–Co, and Al–Cu LDHs. (D–F) X-ray diffraction (XRD) patterns of the LDH formulations showing characteristic reflections assigned to the (003), (006), (009), (015), (018), and (110) planes. (G–I) TEM micrographs of the LDH nanoparticles. Scale bar: 200 nm.

### 3.2 In vitro biological assessment of LDHs

#### 3.2.1 LDH formulations exhibit composition-dependent cytotoxicity in mammalian cell lines

To assess the cytotoxic profile of LDH formulations, cell viability was evaluated using complementary metabolic- and imaging-based assays. MTT dose–response curves were used to assess metabolic activity **(Figure 2)**, while representative merged fluorescence images from the Calcein-AM/Hoechst–PI assay are shown in Figure 3. This imaging-based approach enabled visualization of viable esterase-active cells together with membrane-compromised cells after LDH exposure. To improve visualization clarity in the main manuscript, Figure 3 presents only merged images for each condition, whereas the corresponding individual fluorescence channels are provided in the Supplementary Information for NIH/3T3 **(Figure SI-7)**, HaCaT cells **(Figure SI-8),** A549 **(Figure SI-9)**, and HT-29 **(Figure SI-10)**. Hoechst staining was included to support nuclear segmentation during image analysis; however, in HaCaT cells, acridine orange staining alone provided sufficient nuclear contrast, and the Hoechst channel is therefore not shown separately. Across the tested cell lines, LDH formulations reduced cell viability in a dose-dependent manner, with responses varying according to both metal composition and cellular model **(Supplementary Table S1)**. Overall, Al–Cu showed the strongest cytotoxic profile, including the lowest IC_50_ values in most cell lines, including 47.76 µg/mL in A549 cells in the MTT assay and particularly low IC_50_ values in A549 and HT-29 cells in the Calcein-AM assay. By contrast, Al–Ni generally displayed the highest IC_50_ values, although distinct cell-type-specific responses were observed, such as the pronounced sensitivity of HT-29 cells to this formulation in the MTT assay (IC_50_ = 16.19 µg/mL). The Hoechst–PI assay revealed a more selective pattern, in which Al–Cu remained the most cytotoxic formulation in the human cell lines, whereas Al–Co and Al–Ni caused little or no significant reduction in viability under the same conditions. Together, these results indicate that LDH cytotoxicity is influenced by both metal composition and cell-type context. The comparison across metabolic, fluorescence-based, and imaging-based readouts further 13 suggests that assay modality affects the apparent magnitude of cytotoxicity, reinforcing the value of using orthogonal endpoints for LDH nanosafety assessment.

**Figure 2.**
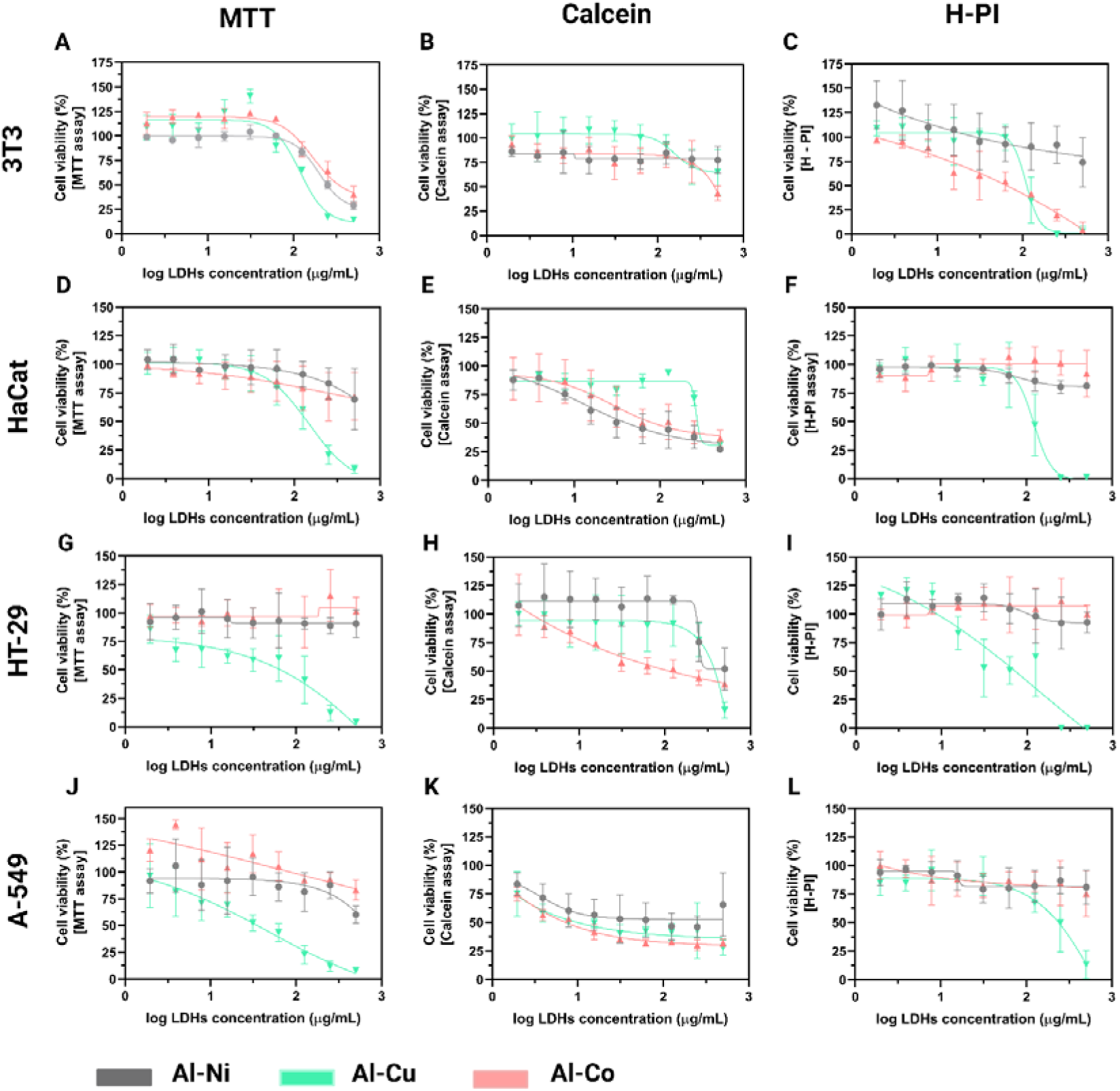
Cytotoxicity profile of Al–Ni, Al–Co, and Al–Cu LDH nanoparticles in mammalian cell lines. Cell viability was assessed after 24 h exposure to increasing nominal LDH concentrations using MTT, Calcein-AM, and Hoechst–PI assays. Panels show viability responses in NIH/3T3 (A–C), HaCaT (D–F), HT-29 (G–I), and A549 (J–L) cells. For each cell line, MTT results are shown in the first column, Calcein-AM results in the second column, and Hoechst–PI imaging-based viability in the third column. Viability values were normalized to untreated controls and plotted against log-transformed nominal LDH concentration (µg/mL). Data are presented as mean ± SD from three independent biological experiments, with three technical replicate wells per condition. Dose–response curves were fitted using nonlinear regression to estimate IC50 values, which are summarized Supplementary Table S1.

**Figure 3.**
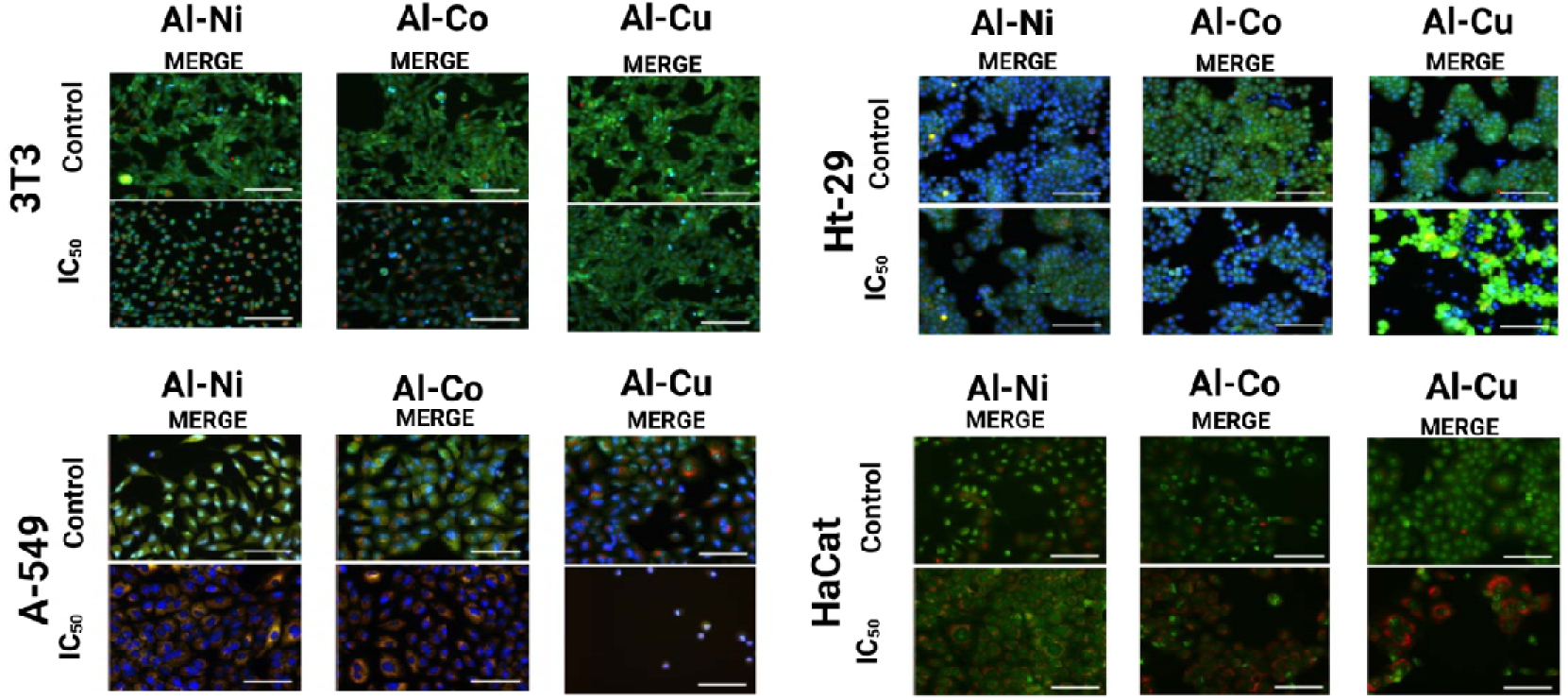
Representative fluorescence images of LDH-induced cytotoxicity in mammalian cell lines. NIH/3T3, HT-29, A549, and HaCaT cells were exposed for 24 h to Al–Ni, Al–Co, or Al–Cu LDHs at the corresponding IC50 concentration. NIH/3T3, HT-29, and A549 cells were stained with Hoechst to identify improved nuclei segmentation, and all cells were stained with Calcein-AM to visualize esterase-active viable cells, and propidium iodide (PI) to identify membrane-compromised cells. Images show the merged overlays. Images were acquired using a 20× objective. Scale bars: 100μm.

#### 3.2.2 High-content imaging discriminates composition-dependent LDH phenotypic signatures and detects subtle low-dose cellular responses

To extend the conventional cytotoxicity assays and capture phenotypic changes beyond bulk viability measurements, high-content imaging (HCI) was performed using the Live Cell Painting method. Single-cell multiparametric profiles were generated for all cell lines following 24h exposure to Al–Ni, Al–Co, and Al–Cu LDHs at IC_50_, IC_50/10_, and IC_50/100_, enabling the assessment of both pronounced and subtle cellular responses **(Figures SI-2 to S-5).** Image-derived features were aggregated into well-level phenotypic profiles and analyzed by linear discriminant analysis (LDA), followed by k-means clustering to identify treatment-associated phenotypic groupings (**Figure 4 and SI-6**).

**Figure 4.**
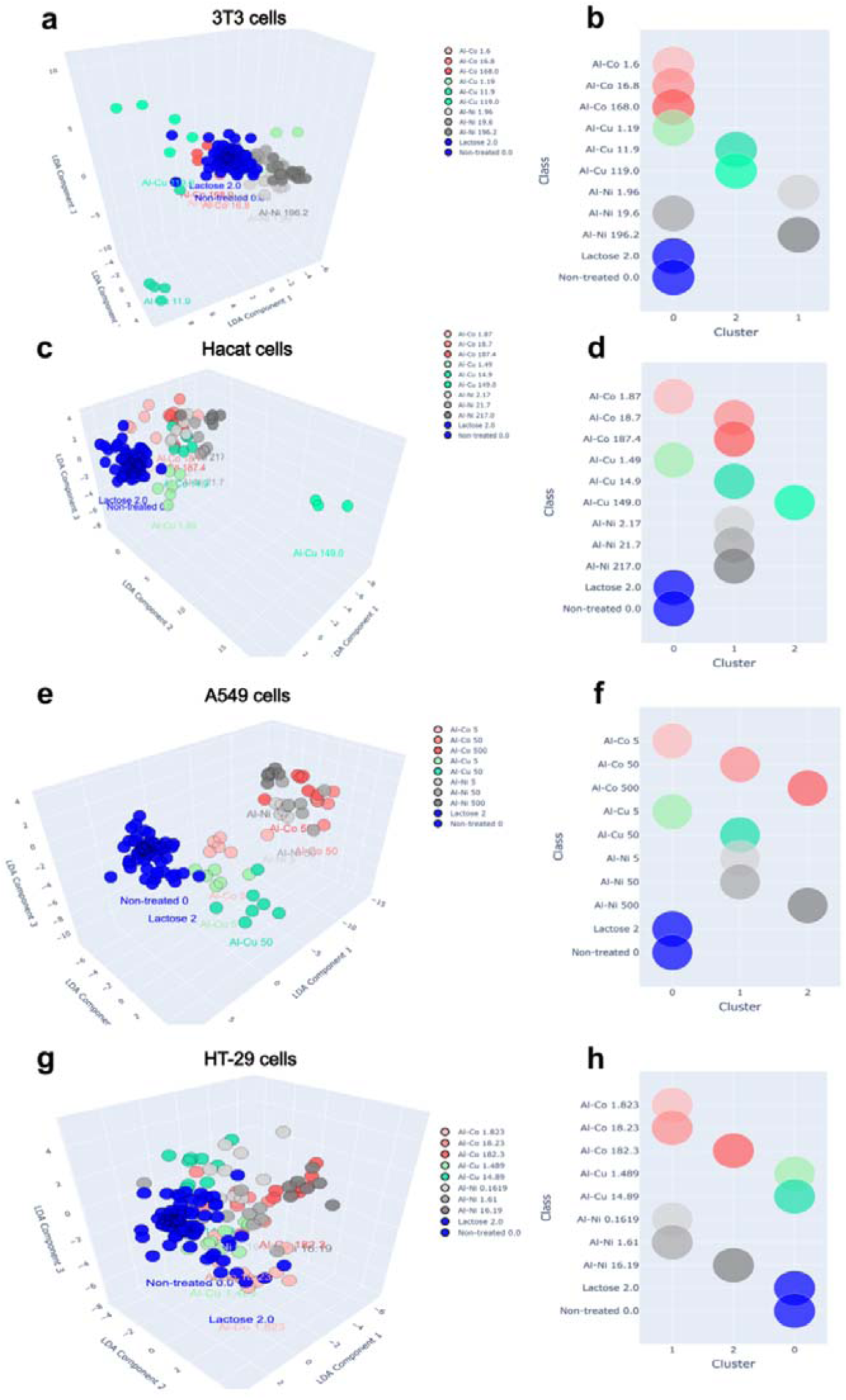
High-content phenotypic profiling of LDH-treated mammalian cells. NIH/3T3, HaCaT, A549, and HT-29 cells were exposed for 24 h to Al–Ni, Al–Co, or Al–Cu LDHs at IC50, IC50/10, and IC50/100 concentrations. Cells were stained using the Live Cell Painting assay and imaged by high-content fluorescence microscopy. Single-cell features were extracted, aggregated into well-level profiles, normalized to negative controls, and analyzed by LDA. Panels A, C, E, and G show 3D LDA projections for NIH/3T3, HaCaT, A549, and HT-29 cells, respectively. Panels B, D, F, and H show the corresponding LDA cluster assignments for each treatment condition. Phenotypic profiles were generated from three technical replicate wells per condition across three independent biological experiments.

Across formulations, IC_50_ exposure induced visible phenotypic alterations, including increased acridine orange signal, changes in acidic vesicle organization, and altered nuclear morphology (**Figure 3 and SI-6).** Among the tested formulations, Al–Cu produced the most pronounced phenotypic shifts; for several cell lines, the highest Al–Cu concentrations led to extensive cell death and debris, resulting in their exclusion during the image quality control stage. In contrast, Al–Ni induced comparatively milder responses. The multivariate analysis supported these visual observations. Untreated and lactose-treated cells clustered closely in all cell lines, supporting assay stability and indicating that the observed separations were treatment-associated rather than driven by the reference condition (**Figure 4**). At IC_50,_ LDH treatments showed marked phenotypic separation from controls, with major contributing features related to cell and compartment morphology, nucleoli and vesicle organization, nuclear intensity distribution, and cytoplasmic granularity (**Figure SI-6**). Notably, Al–Cu exhibited phenotypic alterations starting at the intermediate concentration (IC_50_/10) in 3T3, HaCaT, and A549 cells, where treated samples already appeared phenotypically distinct from corresponding controls. At the highest concentration, data for A549 and HT-29 cells are not shown, as the extensive damage induced was sufficient for these conditions to be excluded during image quality control. Al–Ni nanoparticles also demonstrated pronounced phenotypic activity, showing clear separation from controls across nearly all IC_50_ concentrations in all evaluated cell lines (**Figure 4**), indicating that this formulation also induces measurable cellular perturbations. By contrast, Al–Co produced the weakest overall phenotypic effects, with no evident phenotypic change in NIH/3T3 cells and only limited separation in HT-29 cells at IC_50_. HT-29 cells were a notable exception, as all three Al–Co concentrations showed phenotypic activity; however, these differences do not necessarily indicate increased cytotoxicity. High-content imaging was more sensitive than the conventional viability assays, as composition-dependent phenotypic signatures remained detectable at IC_50/10_ and even IC_50/100_ concentrations that produced little or no measurable effect in the MTT, Calcein-AM, or Hoechst–PI assays. Collectively, HCI detected treatment-associated phenotypic deviations at concentrations that produced little or no measurable effect in conventional MTT, Calcein-AM, or Hoechst–PI assays, supporting its value for identifying sublethal cellular perturbations induced by LDH nanoparticles.

#### 3.2.3 LDH nanoparticles induce composition- and cell type-dependent genotoxic, proliferative, and morphometric changes

Immunocytochemical and morphometric analyses were performed to further assess cellular stress responses induced by Al–Ni, Al–Co, and Al–Cu LDHs. The combined evaluation of γ-H2AX, Ki-67, and phalloidin staining enabled the assessment of DNA damage-associated signaling, proliferation-related status, and cytoskeletal/morphological alterations, respectively^45,46^ **(Figure 5)**. Representative images in Figure 5 show the Al–Cu formulation, which produced the most pronounced biological effects across the evaluated cell lines. Corresponding data for Al–Ni and Al–Co are provided in the Supplementary Information **(Figures SI-7 to SI-10)**. In NIH/3T3 cells, γ-H2AX fluorescence intensity increased with increasing nanoparticle concentration, with the most pronounced response observed after LDH-Al–Cu treatment **(Figure 5A)**. At 250 µg/mL, a reduction in γ-H2AX intensity was detected **(Figure 5A-I)**. In parallel, an increase in Ki-67 intensity was observed at higher nominal concentrations. Morphometric analysis revealed a significant reduction in the “Area Shape” parameter at 250 µg/mL, accompanied by cellular retraction and rounding, as evidenced by phalloidin staining **(Figure 5A-II)**. In A549 cells, γ-H2AX intensity increased at intermediate concentrations, particularly for the Al–Cu formulation **(Figure 5B)**. Ki-67 expression remained stable at lower concentrations, with an increase detected only at the highest nominal exposure concentration of Al–Cu. Morphometric analysis revealed increased cellular heterogeneity, characterized by the coexistence of retracted cells and cells with elongated morphology **(Figure 5B-II)**. In HaCaT cells, γ-H2AX induction was mild across formulations and increased mainly at the highest concentration, without statistically significant differences **(Figure 5C)**. Ki-67 expression remained relatively stable. Phalloidin images indicated partial loss of actin filament organization, particularly in samples treated with LDH-Al–Cu, along with a reduction in the “Area Shape” parameter, which was more pronounced at 250 µg/mL **(Figure 5C-II)**. In HT-29 cells, γ-H2AX staining increased across the tested formulations, including at intermediate nominal exposure concentrations **(Figure 5D)**. Concomitantly, a marked reduction in Ki-67 expression and pronounced disorganization of actin filaments were observed, along with cellular retraction and loss of cytoplasmic definition. These effects were quantitatively reflected by a significant decrease in the “Area Shape” parameter **(Figure 5D-II)**. Overall, LDHs induced distinct responses between non-tumor cells (NIH/3T3 and HaCaT) and tumor cells (A549 and HT-29). LDH-Al–Cu elicited the most pronounced effects, followed by LDH-Al–Ni, whereas LDH-Al–Co displayed an intermediate behavior, remaining closer to control conditions. Overall, these data indicate that LDH exposure induces composition- and cell type-dependent stress-associated and morphometric changes. Al–Cu produced the strongest overall response, whereas Al–Co remained closer to control conditions in most models. These findings support the cytotoxicity and HCI results by showing that LDH exposure is associated with DNA damage-related signaling, altered cytoskeletal organization, and changes in cell morphology.

**Figure 5.**
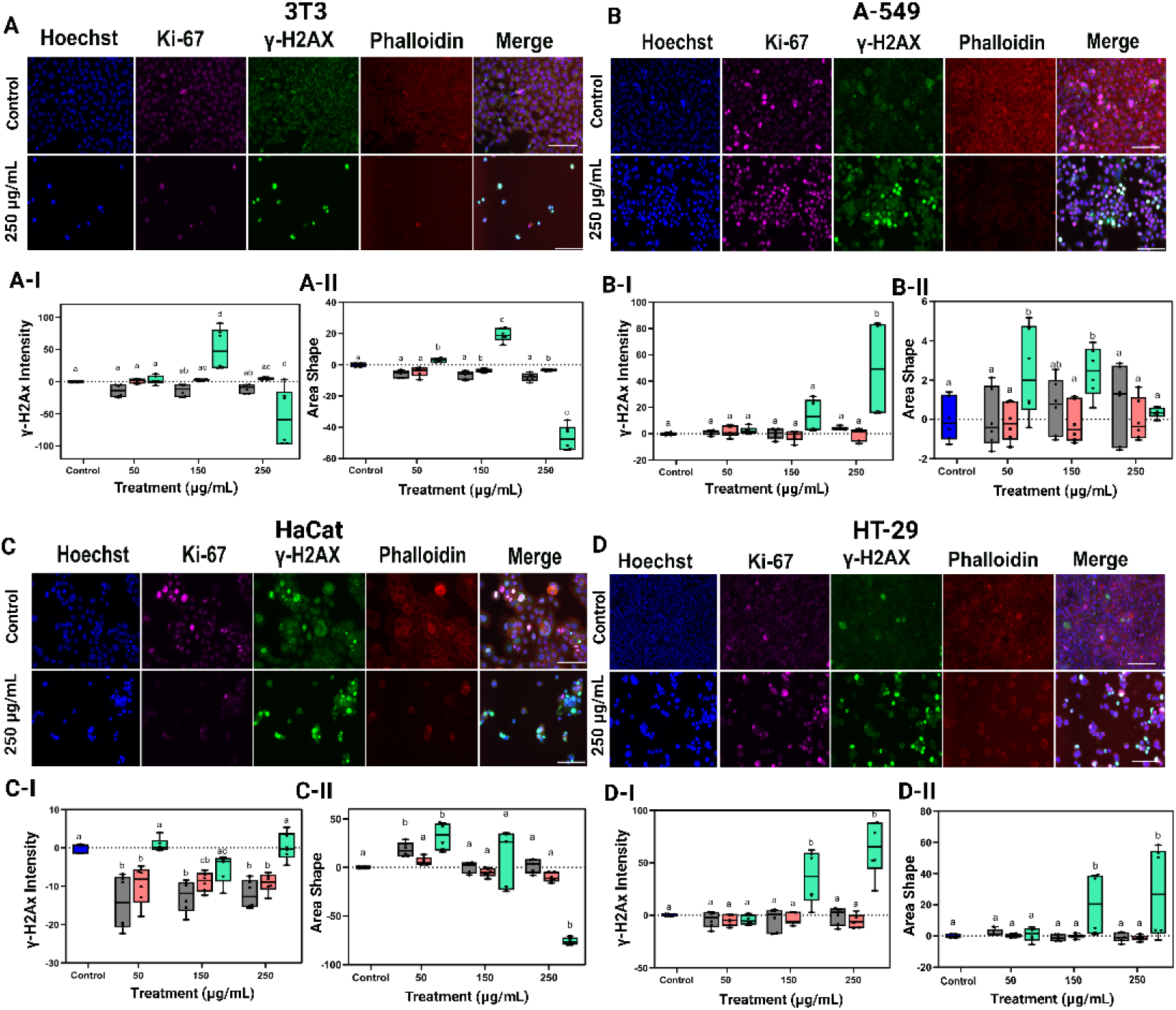
Immunocytochemical and morphometric analysis of LDH-treated mammalian cells. NIH/3T3 (A), A549 (B), HaCaT (C), and HT-29 (D) cells were exposed to LDH nanoparticles and stained with Hoechst to identify nuclei, Ki-67 as a proliferation-associated marker, γ-H2AX as a DNA damage-associated marker, and phalloidin to visualize the actin cytoskeleton. Representative fluorescence images are shown for Al–Cu-treated cells. Lower panels show quantification of γ-H2AX fluorescence intensity (I) and the Area Shape morphometric parameter (II) after treatment at the indicated concentrations. Data are presented as mean ± SD from three independent biological experiments. Statistical analysis was performed using two-way ANOVA followed by Tukey’s post hoc test, with p < 0.05 considered significant. Scale bar 100 μm.

## 4. Discussion

In this study, we tested the hypothesis that the biological effects of LDH nanoparticles are composition-dependent by comparing Al–Ni, Al–Co, and Al–Cu formulations across cell models relevant to inhalation, ingestion, and dermal exposure, together with a comparative murine fibroblast model. To examine this, we combined conventional viability assays (MTT, Calcein-AM, and fluorescence-based cell counting) with immunocytochemical, morphometric, and high-content imaging approaches, including Live Cell Painting. This integrated strategy allowed us to capture not only overt cytotoxicity but also genotoxicity-associated responses and subtle phenotypic alterations across biologically relevant models. Overall, the data support a clear composition-dependent toxicity profile for LDHs and provide an experimental basis for the safer development and risk assessment of LDH-based technologies in agricultural and related applications.

The central goal of this study was to determine how the metal composition of LDH nanoparticles influences their cytotoxic profile in mammalian cells. This question is directly relevant to the development of safer and more effective nanoparticle-based technologies. By combining conventional viability assays (MTT, Calcein-AM, and fluorescence-based cell counting) with immunocytochemical, morphometric, and high-content imaging approaches, including Live Cell Painting, we found that LDH toxicity was strongly composition-dependent, consistent with the broader principle that nanomaterial composition is a major determinant of biological response^47,48^. Overall, Al–Cu was the most biologically disruptive formulation, causing the greatest reduction in cell viability across the conventional assays, pronounced changes in cell morphology, and stronger alterations in γ-H2AX- and Ki-67-associated responses, in line with previous reports describing the higher biological reactivity of copper-containing nanomaterials^49–51^. In addition, Al–Cu induced clear phenotypic signatures even at concentrations as low as IC_50/100_, indicating that its effects extend beyond overt cytotoxicity. These findings suggest that even sublethal exposures are sufficient to trigger relevant cellular responses, including early structural and functional alterations. Al–Ni showed intermediate activity, with measurable cytotoxicity in the viability assays and more limited phenotypic and morphometric alterations. The lower magnitude of the observed effects suggests that, although biologically active, Al–Ni may present a less aggressive cellular interaction profile compared to Al–Cu, possibly reflecting differences in surface reactivity and cellular uptake dynamics. By contrast, Al–Co was the mildest formulation overall, producing weaker cytotoxic effects and comparatively subtle phenotypic changes; however, specific cell lines such as HaCaT and HT-29 exhibited distinct sensitivity patterns, likely reflecting cell-type-dependent differences in nanoparticle uptake and stress response pathways^52^. Together, these findings show that LDH toxicity depends not only on formulation composition but also on cellular context, reinforcing the importance of composition-aware safety assessment during nanomaterial development.

Another important aspect of nanoparticle toxicology is the extent to which the nanoform itself contributes to the observed biological response^53^. The LDH nanoparticles evaluated here showed high colloidal and thermal stability, relatively homogeneous dispersions, and preservation of the characteristic lamellar structure, all of which are desirable features for technological applications. These properties are especially relevant for applications requiring robustness under variable environmental conditions, including agriculture^19,54,55^. Nevertheless, none of the physicochemical characteristics measured for the Al–Cu formulation was sufficient to explain its substantially greater cytotoxicity. This suggests that the relatively modest physicochemical differences among the selected formulations cannot, by themselves, explain the marked divergence in biological effects. Rather, LDH composition appears to be a major factor shaping the biological response, although differences in nanoparticle uptake, intracellular trafficking, dissolution, and metal ion release may also contribute and should be investigated in future studies^56^. In this sense, our results are compatible with redox- and dissolution-based models of nanoparticle toxicity, in which differences in intracellular reactivity lead to distinct biological responses despite similar bulk physicochemical properties^57^. The stronger activity of Al–Cu and the comparatively mild behavior of Al–Ni therefore support the view that the divalent metal within the LDH lattice is a major determinant of biological response.

Our results also highlight the sublethal effects of LDH formulations in mammalian cells. Such effects are particularly relevant because they may better approximate realistic exposure scenarios than endpoints based solely on overt loss of viability^58,59^. In this study, increased γ-H2AX signal indicated DNA damage-associated stress, consistent with the use of γ-H2AX as a sensitive marker of double-strand break responses and genotoxic stress in nanoparticle-exposed cells^46,60^. This response was especially evident after exposure to the Al–Cu formulation, supporting the view that this composition triggers stronger intracellular stress than the other LDHs tested. In parallel, phalloidin staining and morphometric analysis revealed cytoskeletal remodeling, cell retraction, and altered spreading, features that are consistent with the known ability of nanomaterials to disturb actin organization during uptake and stress adaptation^61,62^. By contrast, Ki-67 levels showed only limited variation. Given that Ki-67 is a proliferation-associated marker with functions extending beyond a simple binary indicator of cell division, these results suggest that LDH exposure did not markedly alter proliferation-associated status under the conditions evaluated, even when other sublethal stress responses were already detectable^63,64^. Taken together, these data suggest that sublethal LDH exposure, particularly for Al–Cu, is characterized less by an immediate decrease of proliferation and more by early stress responses involving DNA damage-associated signaling and cytoskeletal remodeling.

Image-based phenotypic profiling captures rich information at the cellular and subcellular levels and therefore provides a broader view of cell health and homeostasis than bulk viability measurements alone^65^. In this study, high-content imaging (HCI) identified formulation-specific phenotypes that remained detectable at concentrations below those affecting conventional viability assays. Using the Live Cell Painting approach, we were able to resolve subtle cellular alterations that would have been overlooked by standard cytotoxicity readouts. We recently reported a similar increase in sensitivity in the evaluation of silver nanoparticles, reinforcing the value of HCI for detecting early or sublethal nanoparticle-induced perturbations^66,67^. This is particularly important for toxicological assessment at low exposure levels, which may better reflect realistic exposure scenarios. By extending viability analysis beyond a simple “alive versus dead” classification, Live Cell Painting enabled the identification of multiparametric single-cell phenotypes, including changes in nuclear organization and acidic vesicle patterning, that were invisible to conventional endpoints but informative for mechanistic interpretation. Taken together, these findings indicate that combining orthogonal viability assays with high-content imaging improves both sensitivity and biological interpretability, allowing cytotoxicity to be assessed in an early phenotypic window that precedes overt cell death.

Although *in vitro* models cannot fully reproduce the complexity of *in vivo* exposure, the four mammalian cell lines used in this study revealed both common toxicity patterns and cell type-specific responses. An interesting example is the behavior of the Al–Ni formulation, which showed distinctive activity in the skin-related models, particularly HaCaT cells. While this observation should be interpreted cautiously, it is noteworthy in light of the known association between nickel exposure and allergic contact dermatitis in humans^68^. These findings are in agreement with the literature that suggests that keratinocytes, including HaCaT, can initiate and help shaping the inflammatory response triggered by nickel^69,70^. In this sense, the present findings may help identify biologically relevant response patterns that can serve as starting points for future *in vivo* toxicity studies. Further advances in understanding LDH toxicity will require deeper investigation of nanoparticle uptake mechanisms, intracellular dissolution, and metal ion release, as these processes may be directly linked to the cytotoxic effects observed here. Combined with more detailed mechanistic follow-up, particularly for the Al–Cu formulation, such studies could help guide the design of safer LDH nanoparticles. Despite these limitations, the present study shows that the safety of LDH nanoparticles cannot be assumed across compositions and highlights the need for composition-aware evaluation during the development of LDH-based technologies.

## 5. Conclusion

This study provides a comparative evaluation of the biological effects of Al–Ni, Al–Co, and Al–Cu layered double hydroxide (LDH) nanoparticles across mammalian cell models representing relevant exposure-associated tissues. All formulations exhibited colloidal stability and preserved lamellar structure, yet their biological responses differed markedly. Conventional cytotoxicity assays established an overall toxicity trend of Al–Cu > Al–Co > Al–Ni, while high-content imaging revealed formulation-specific phenotypic alterations that remained detectable at sublethal concentrations. In particular, Al–Cu induced pronounced vesicular and nuclear perturbations, whereas Al–Ni produced comparatively milder effects across most models. These findings demonstrate that LDH biological activity is strongly composition-dependent and further show that high-content phenotypic profiling provides additional sensitivity for detecting treatment-associated cellular perturbations beyond conventional viability endpoints. Collectively, the comparatively milder biological profile of Al–Ni supports its further investigation as a potentially safer candidate for LDH-based agricultural delivery systems, whereas the stronger activity observed for Al–Cu highlights the importance of composition-aware nanosafety assessment during the development of LDH-based technologies.

## 6. Acknowledgements

This study was funded by the Coordination for the Improvement of Higher Education Personnel – Brazil (CAPES) – Funding Code 001; the São Paulo Research Foundation (FAPESP; grants no. 2023/05204-5, 2023/06143-0, 2022/03250-7, and 2020/01218-3); the National Council for Scientific and Technological Development (CNPq); and the Support Fund for Teaching, Research, and Extension (FAEPEX–UNICAMP; grants no. 2633/17, 2421/20, 2237/21, 2178/22, and 2534/23).

The Article Processing Charge (APC) for this publication was funded by the Coordination for the Improvement of Higher Education Personnel – CAPES (ROR identifier: 00×0ma614). For the purposes of open access, the authors have applied a Creative Commons Attribution (CC BY) license to any accepted version of the article.

