## Supplemental material for "Layered double hydroxide nanoparticles induce composition-dependent cytotoxic and phenotypic effects in mammalian cells"

**Table S1. IC_50_ values of LDH nanoparticles in multiple mammalian cell lines determined using complementary viability assays.** IC_50_ values are expressed in µg/mL and were estimated by nonlinear regression analysis of dose–response curves obtained from MTT, Calcein-AM, and Hoechst/PI (H-PI) assays. Approximate values are indicated by “~”. Entries labeled as “NR” indicate that the IC_50_ was not reached within the tested concentration range, whereas “BTR” indicates that the IC_50_ was below the tested concentration range. “Undetermined” indicates unreliable curve fitting or insufficient response for IC_50_ estimation.

| NIH/3T3 | | | |
| --- | --- | --- | --- |
| Assay | **Al-Ni** | **Al-Co** | **Al-Cu** |
| MTT | 196.2 | 168.3 | 119.2 |
| CALCEIN | $\sim$10.6 | $NR$ | 157.1 |
| H-PI | $BTR$ | $NR$ | 109.2 |
| HaCaT | | | |
| Assay | **Al-Ni** | **Al-Co** | **Al-Cu** |
| MTT | NR | 187.4 | 149.0 |
| CALCEIN | 13.4 | 29.1 | $\sim$264.0 |
| H-PI | 76.9 | $\sim$7.7 | 122.3 |
| HT-29 | | | |
| Assay | **Al-Ni** | **Al-Co** | **Al-Cu** |
| MTT | 16.2 | 182.3 | *NR* |
| CALCEIN | $\sim$245.7 | $BTR$ | $\sim$2.5 |
| H-PI | 99.1 | $\sim$7.3 | 132.4 |
| A549 | | | |
| Assay | **Al-Ni** | **Al-Co** | **Al-Cu** |
| MTT | *NR* | $\sim$174.9 | 47.8 |
| CALCEIN | 3.5 | 0.7 | $BTR$ |
| H-PI | 16.2 | Undetermined | *NR* |

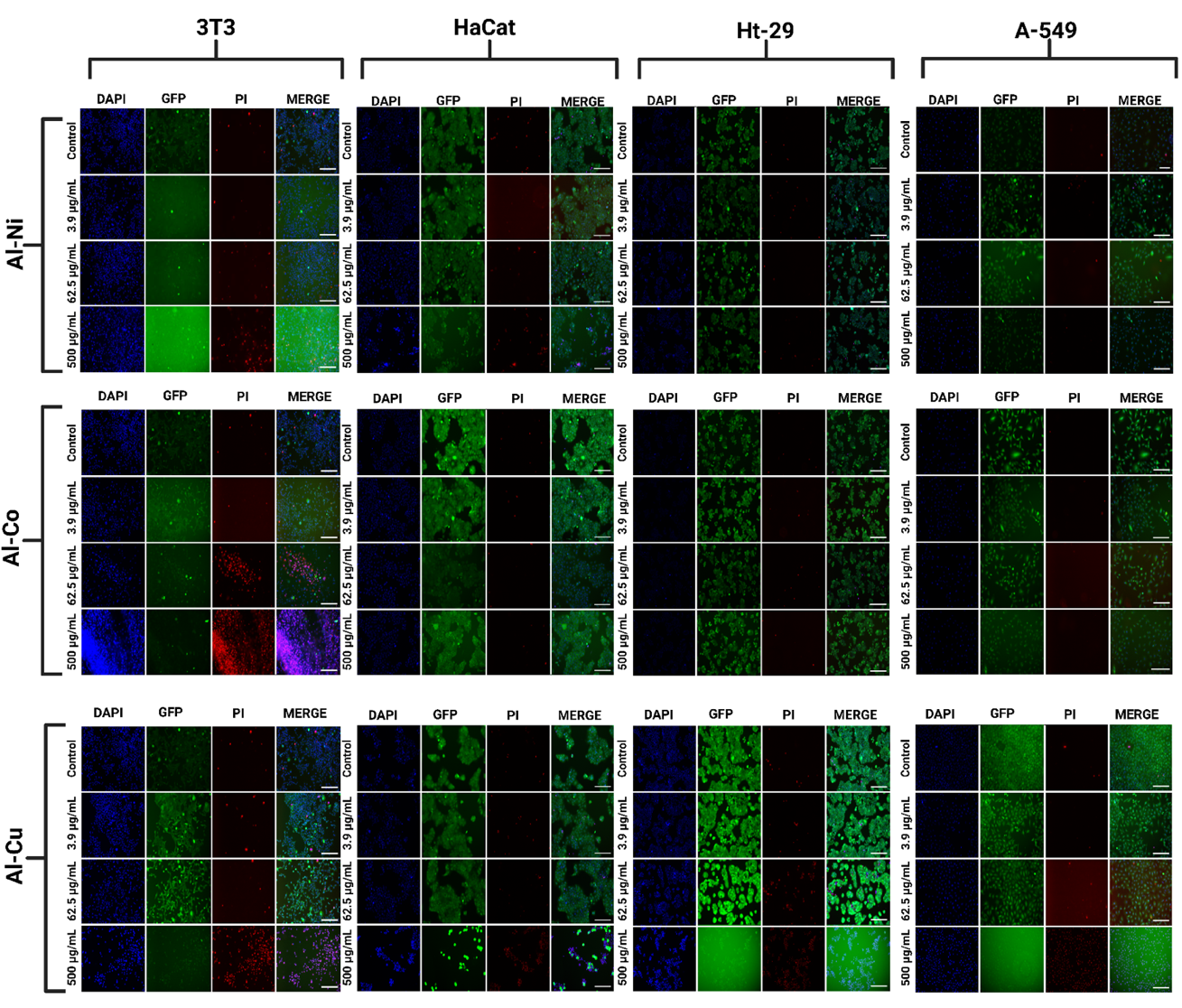

**Figure SI-1. Representative fluorescence images of mammalian cell viability after exposure to LDH nanoparticles**. NIH/3T3, HaCaT, HT-29, and A549 cells were exposed for 24 h to Al–Ni, Al–Co, and Al–Cu layered double hydroxide (LDH) nanoparticles at 3.9, 62.5, and 500 µg/mL, together with untreated controls. Columns show the individual fluorescence channels corresponding to DAPI/Hoechst staining (blue, nuclei), Calcein-AM fluorescence (green, esterase-active viable cells), propidium iodide (PI; red, membrane-compromised cells), and merged images. Rows correspond to increasing LDH concentrations for each formulation. Images were acquired using a 10× objective lens. Scale bars = 200 µm.

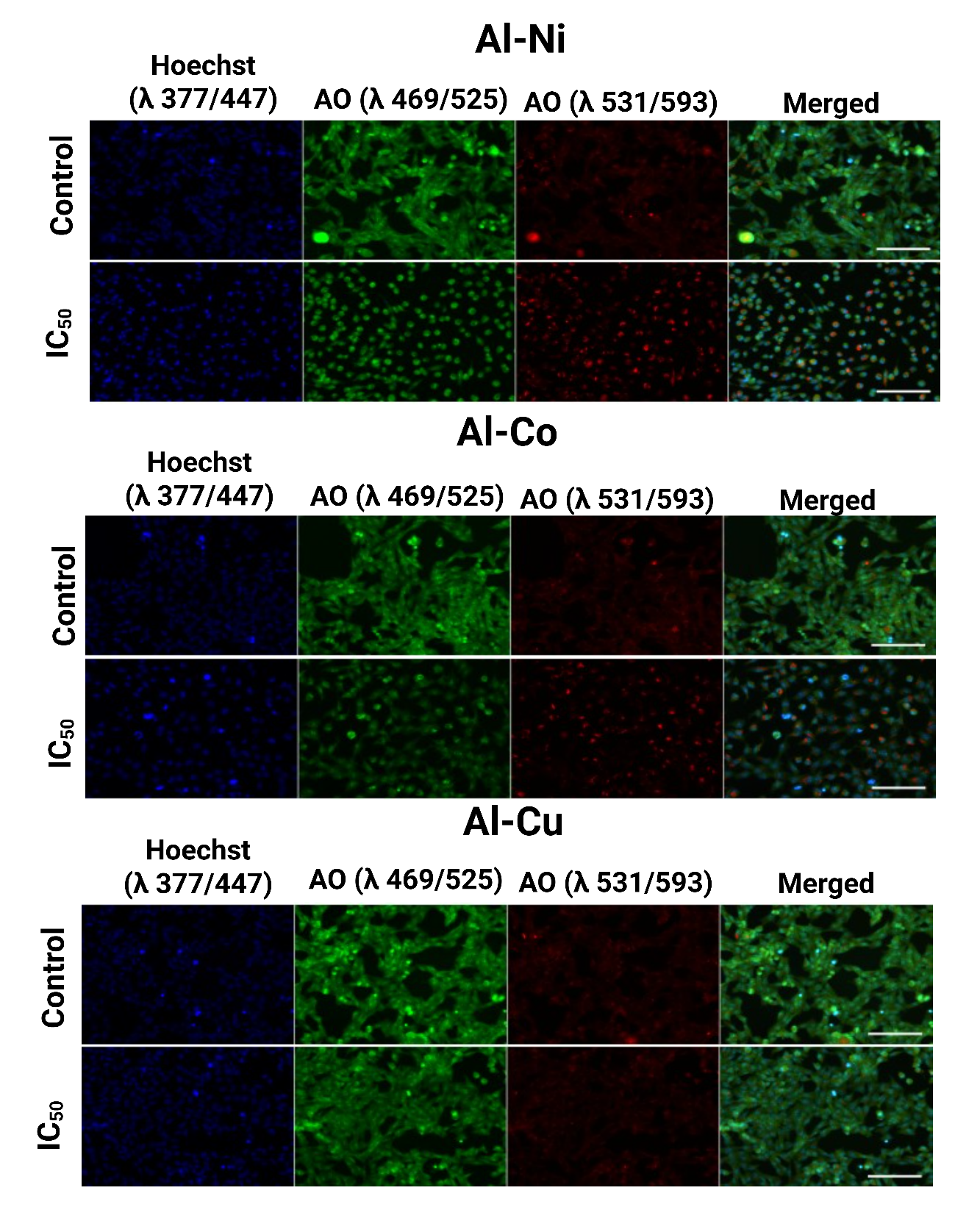

**Figure SI-2. Representative images from the Live Cell Painting assay used to assess phenotypic alterations in NIH/3T3 cells exposed to layered double hydroxide (LDH) nanoparticles.** NIH/3T3 cells were treated for 24 h with Al–Ni, Al–Co, and Al–Cu LDH nanoparticles at the corresponding IC₅₀ concentrations, together with untreated control cells. Columns represent fluorescence channels: Hoechst (DAPI channel, nuclear staining), GFP signal (cytoplasmic, genetic material), PI signal (acidic vesicles / membrane-compromised cells), and a merged image combining all channels. Images were acquired using a 10× objective lens. Scale bar = 200 µm.

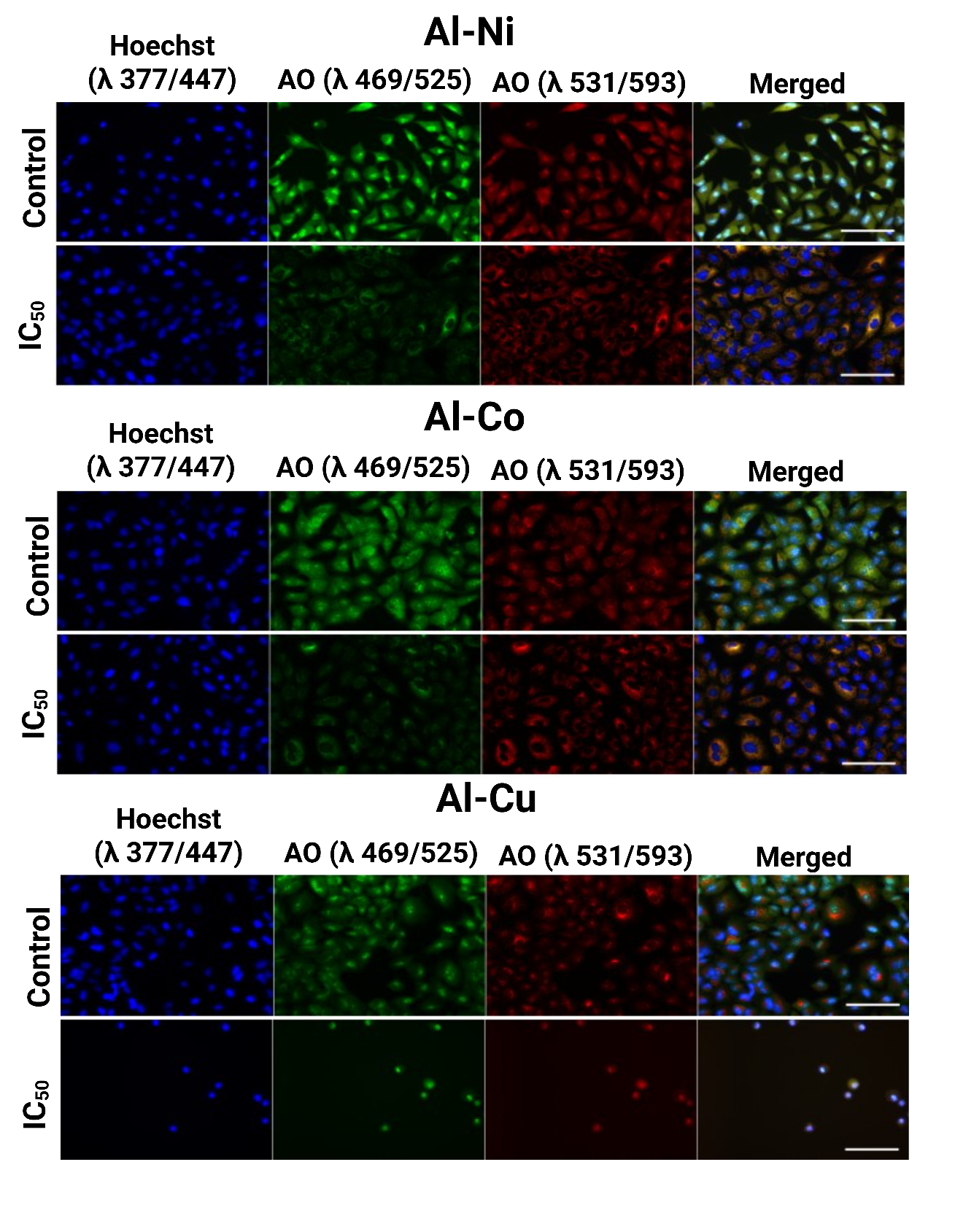

**Figure SI-3. Representative images from the Live Cell Painting assay used to assess phenotypic alterations in A-549 cells exposed to layered double hydroxide (LDH) nanoparticles.** A-549 cells were treated for 24 h with Al–Ni, Al–Co, and Al–Cu LDH nanoparticles at the corresponding IC₅₀ concentrations, together with untreated control cells. Columns represent fluorescence channels: Hoechst (DAPI channel, nuclear staining), GFP signal (cytoplasmic, genetic material), PI signal (acidic vesicles / membrane-compromised cells), and a merged image combining all channels. Images were acquired using a 10× objective lens. Scale bar = 200 µm.

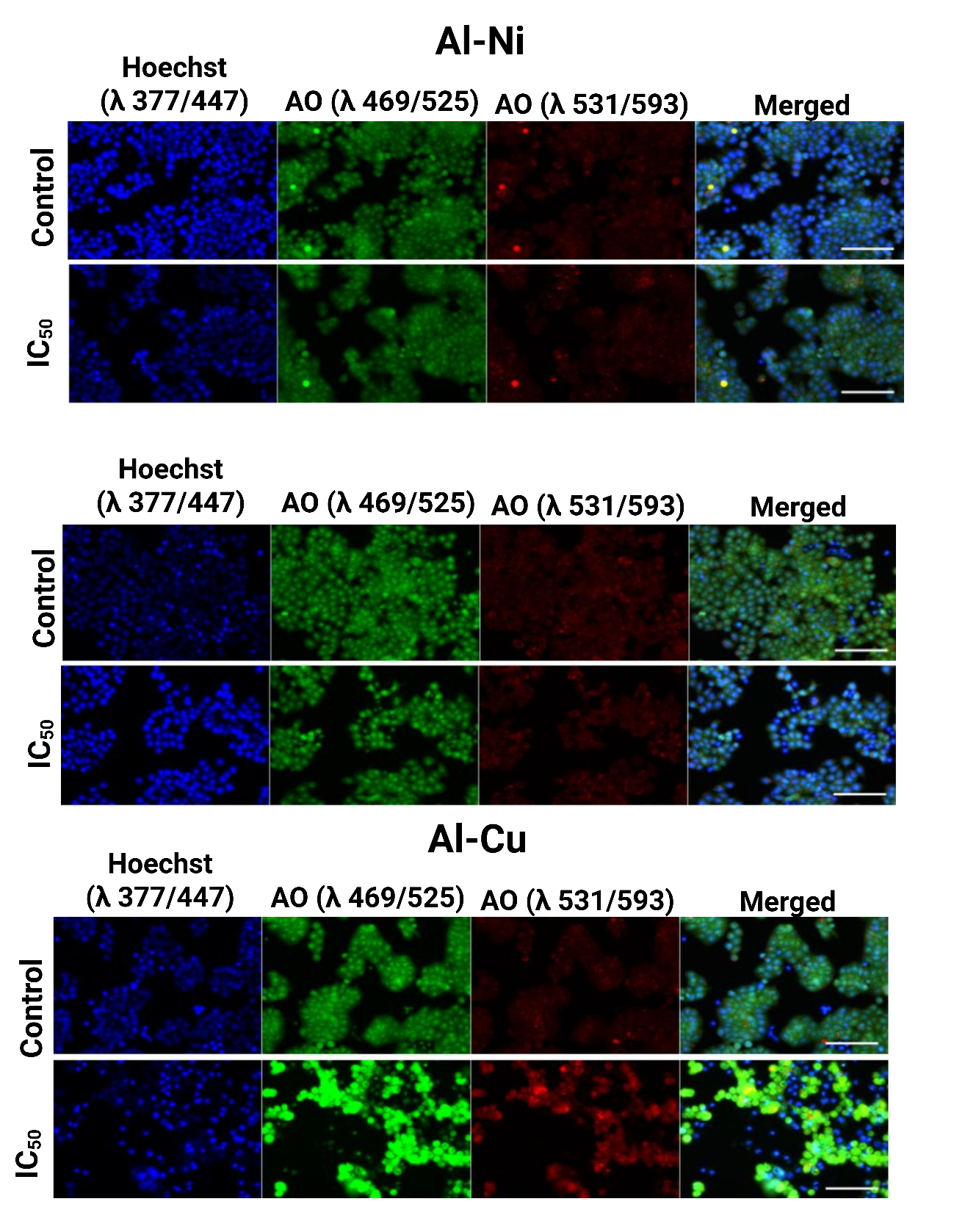

**Figure SI-4. Representative images from the Live Cell Painting assay used to assess phenotypic alterations in HT-29 cells exposed to layered double hydroxide (LDH) nanoparticles.** HT-29 cells were treated for 24 h with Al–Ni, Al–Co, and Al–Cu LDH nanoparticles at the corresponding IC₅₀ concentrations, together with untreated control cells. Columns represent fluorescence channels: Hoechst (DAPI channel, nuclear staining), GFP signal (cytoplasmic, genetic material), PI signal (acidic vesicles / membrane-compromised cells), and a merged image combining all channels. Images were acquired using a 10× objective lens. Scale bar = 200 µm.

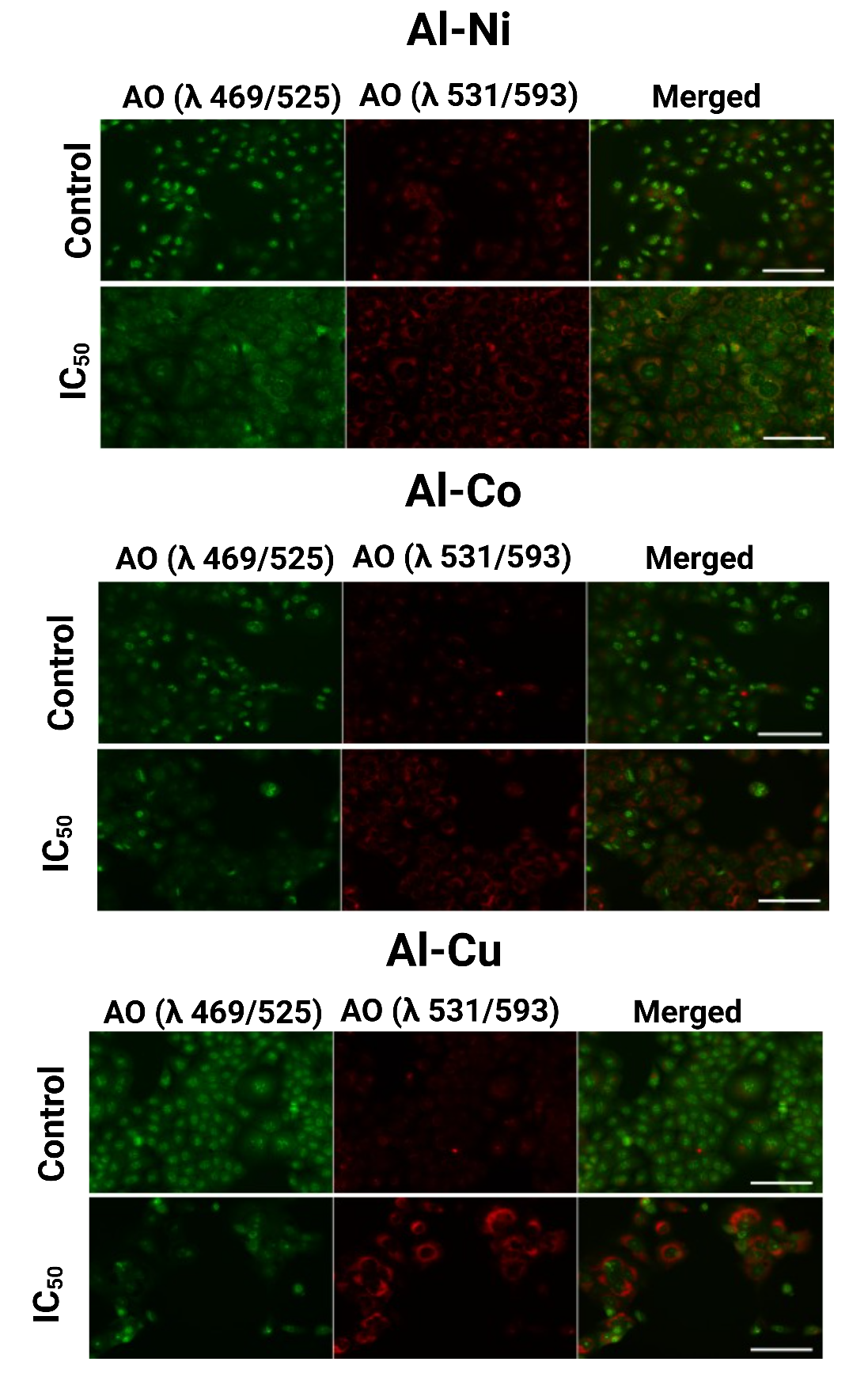

**Figure SI-5. Representative images from the Live Cell Painting assay used to assess phenotypic alterations in HaCat cells exposed to layered double hydroxide (LDH) nanoparticles.** HaCat cells were treated for 24 h with Al–Ni, Al–Co, and Al–Cu LDH nanoparticles at the corresponding IC₅₀ concentrations, together with untreated control cells. Columns represent fluorescence channels: Hoechst (DAPI channel, nuclear staining), GFP signal (cytoplasmic, genetic material), PI signal (acidic vesicles / membrane-compromised cells), and a merged image combining all channels. Images were acquired using a 10× objective lens. Scale bar = 200 µm.

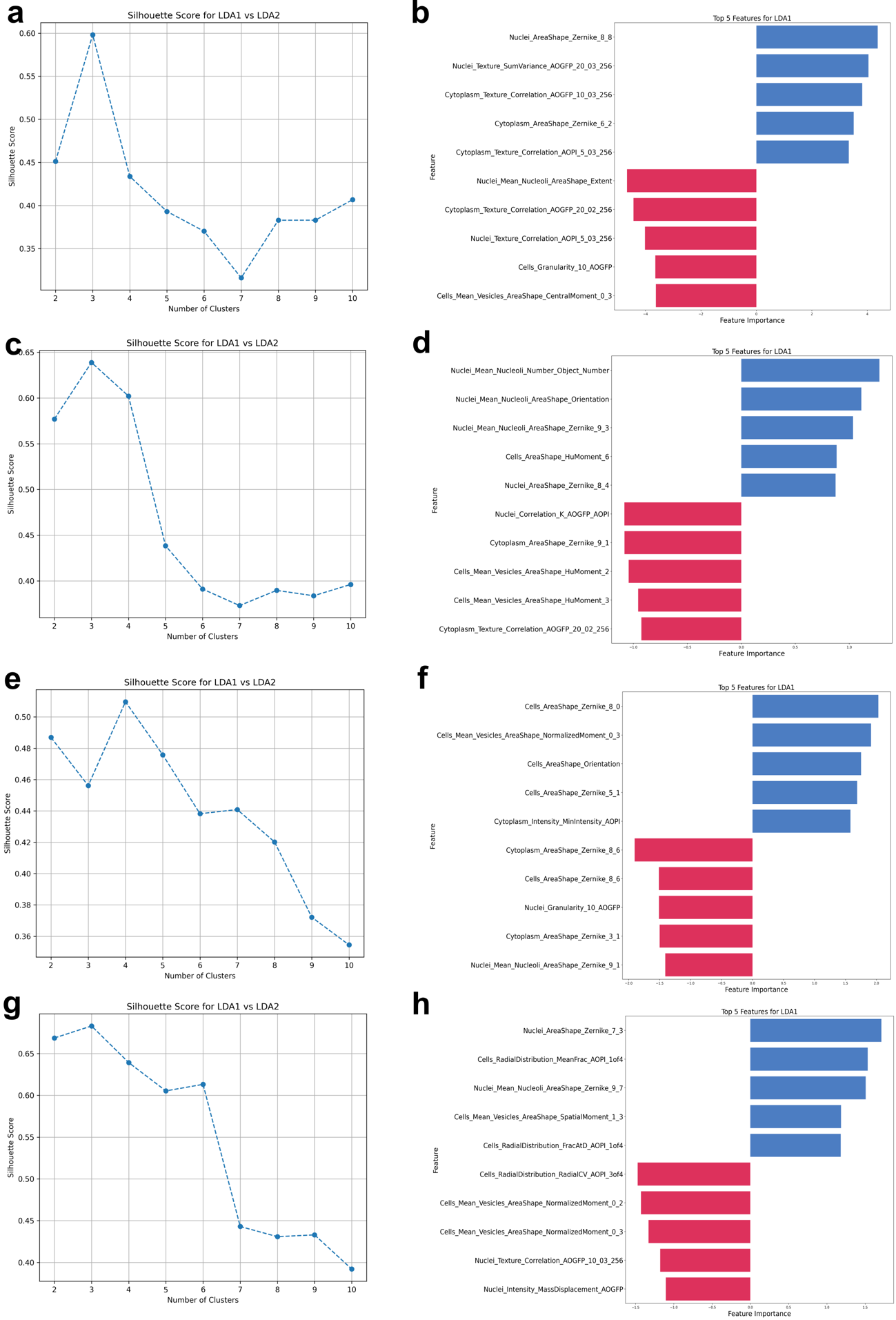

**Figure SI-6. Phenotypic analysis of 3T3, HaCaT, A549, and HT-29 cells. (a, c, e, g) Silhouette plots derived from LDA-transformed data for 3T3, HaCaT, A549, and HT-29 cells, respectively. (b, d, f, h) Top five most important positive and negative Component 1 features contributing to the LDA separation for each cell line.**

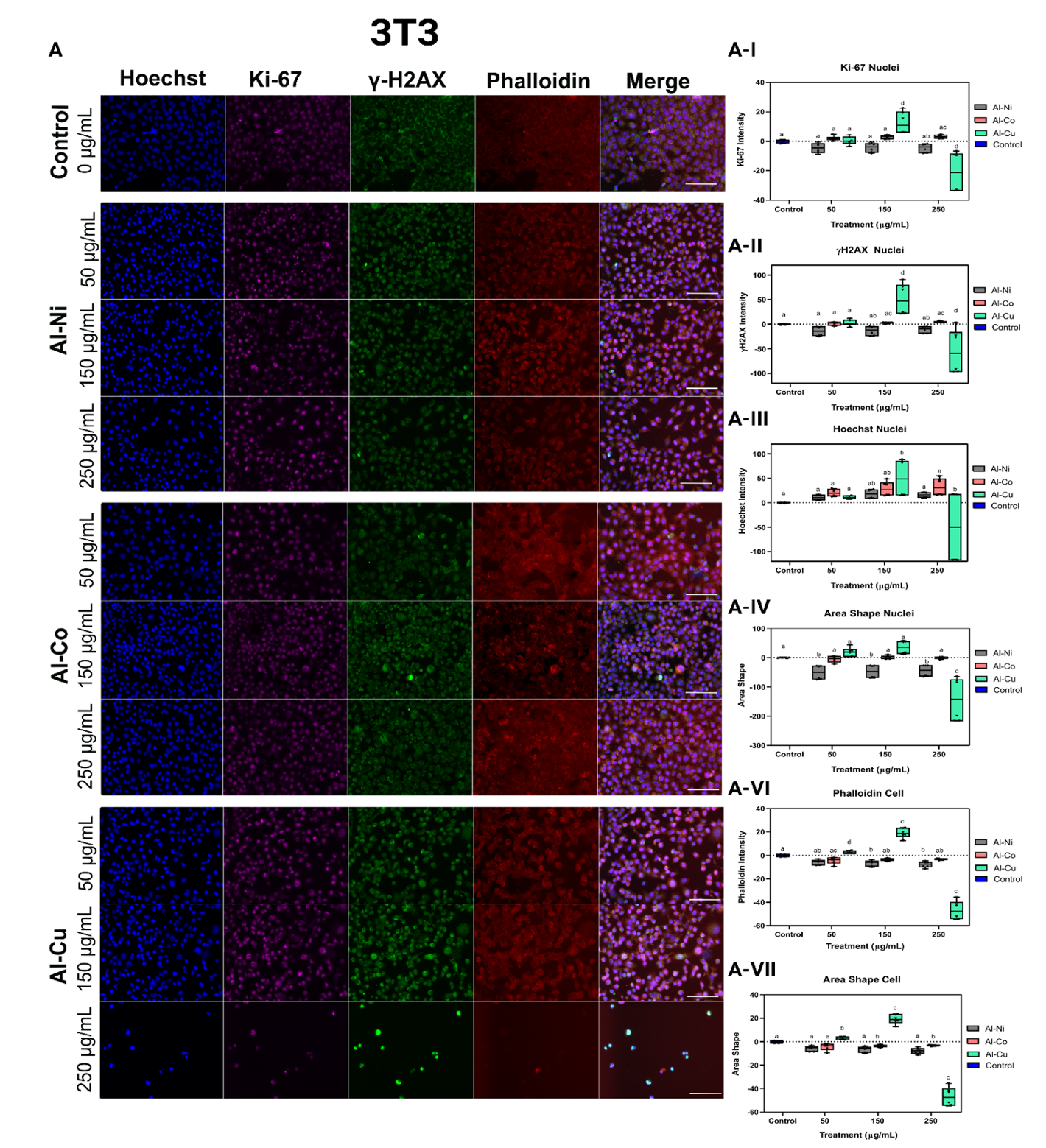

**Figure SI-7. Immunocytochemical and morphometric analysis of NIH/3T3 cells exposed to LDH nanoparticles**. NIH/3T3 cells were treated for 24 h with Al–Ni, Al–Co, and Al–Cu layered double hydroxide (LDH) nanoparticles at 50, 150, and 250 µg/mL. Cells were stained with Hoechst 33342 (blue, nuclei), Ki-67 (magenta, proliferation-associated marker), γ-H2AX (green, DNA damage-associated marker), and phalloidin (red, actin cytoskeleton). Representative fluorescence images are shown together with the corresponding quantitative analyses of fluorescence intensity and morphometric parameters (A-I to A-VII). Scale bar = 100 µm.

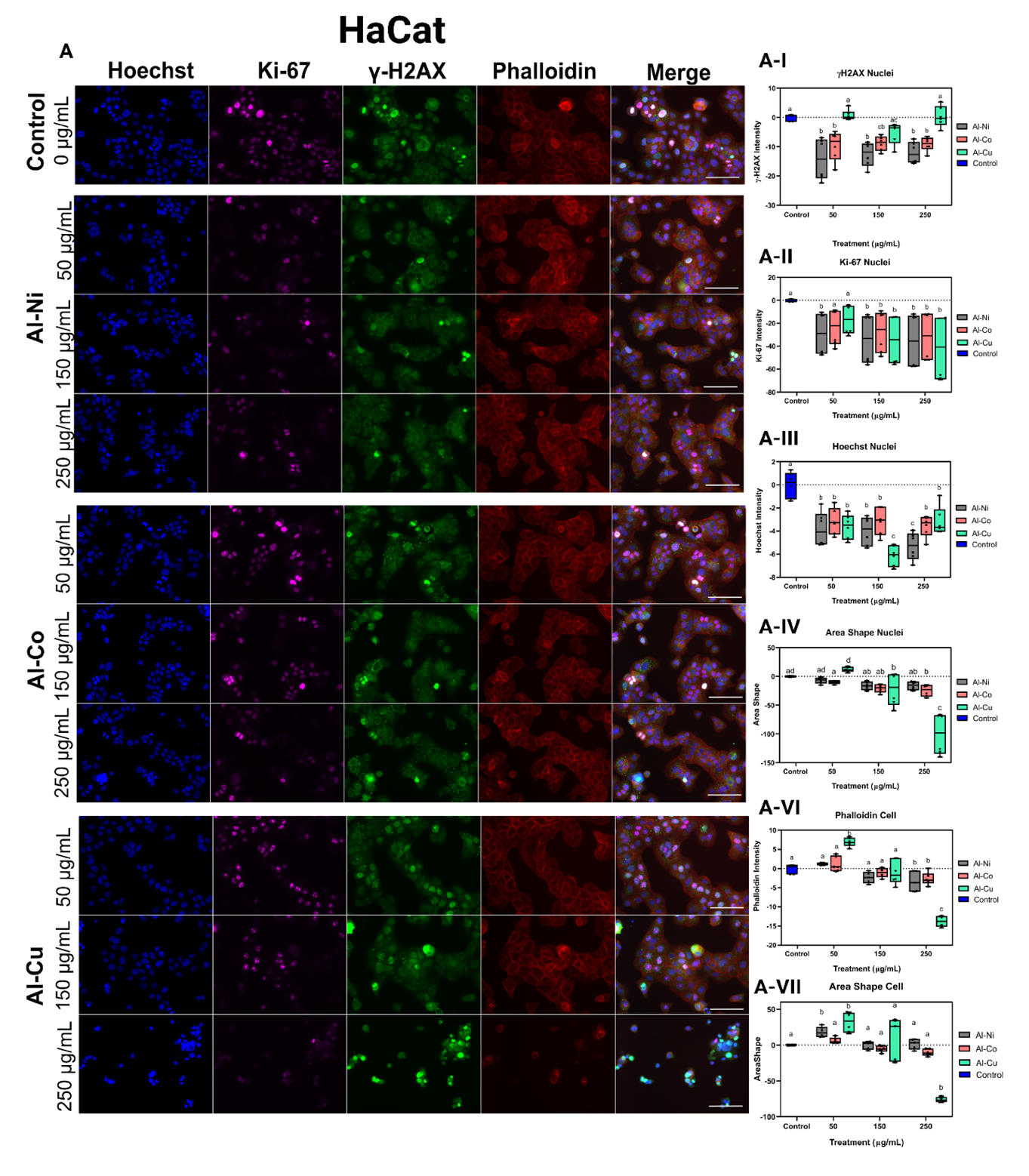

**Figure SI-8. Immunocytochemical and morphometric analysis of HaCaT cells exposed to LDH nanoparticles.** HaCaT cells were treated for 24 h with Al–Ni, Al–Co, and Al–Cu layered double hydroxide (LDH) nanoparticles at 50, 150, and 250 µg/mL. Cells were stained with Hoechst 33342 (blue, nuclei), Ki-67 (magenta, proliferation-associated marker), γ-H2AX (green, DNA damage-associated marker), and phalloidin (red, actin cytoskeleton). Representative fluorescence images are shown together with the corresponding quantitative analyses of fluorescence intensity and morphometric parameters (A-I to A-VII). Scale bar = 100 µm.

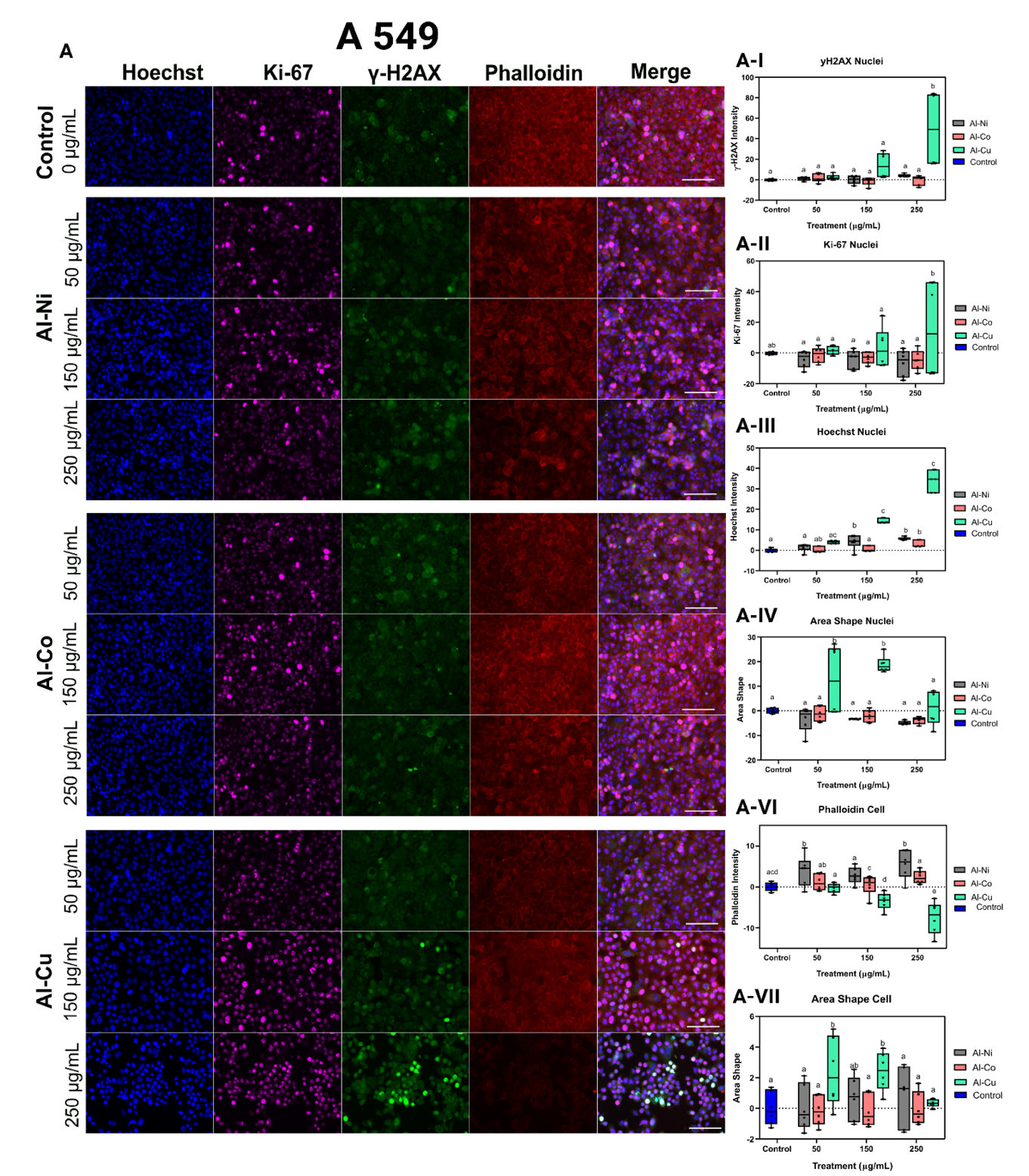

**Figure SI-9. Immunocytochemical and morphometric analysis of A549 cells exposed to LDH nanoparticles.** A549 cells were treated for 24 h with Al–Ni, Al–Co, and Al–Cu layered double hydroxide (LDH) nanoparticles at 50, 150, and 250 µg/mL. Cells were stained with Hoechst 33342 (blue, nuclei), Ki-67 (magenta, proliferation-associated marker), γ-H2AX (green, DNA damage-associated marker), and phalloidin (red, actin cytoskeleton). Representative fluorescence images are shown together with the corresponding quantitative analyses of fluorescence intensity and morphometric parameters (A-I to A-VII). Scale bar = 100 µm.

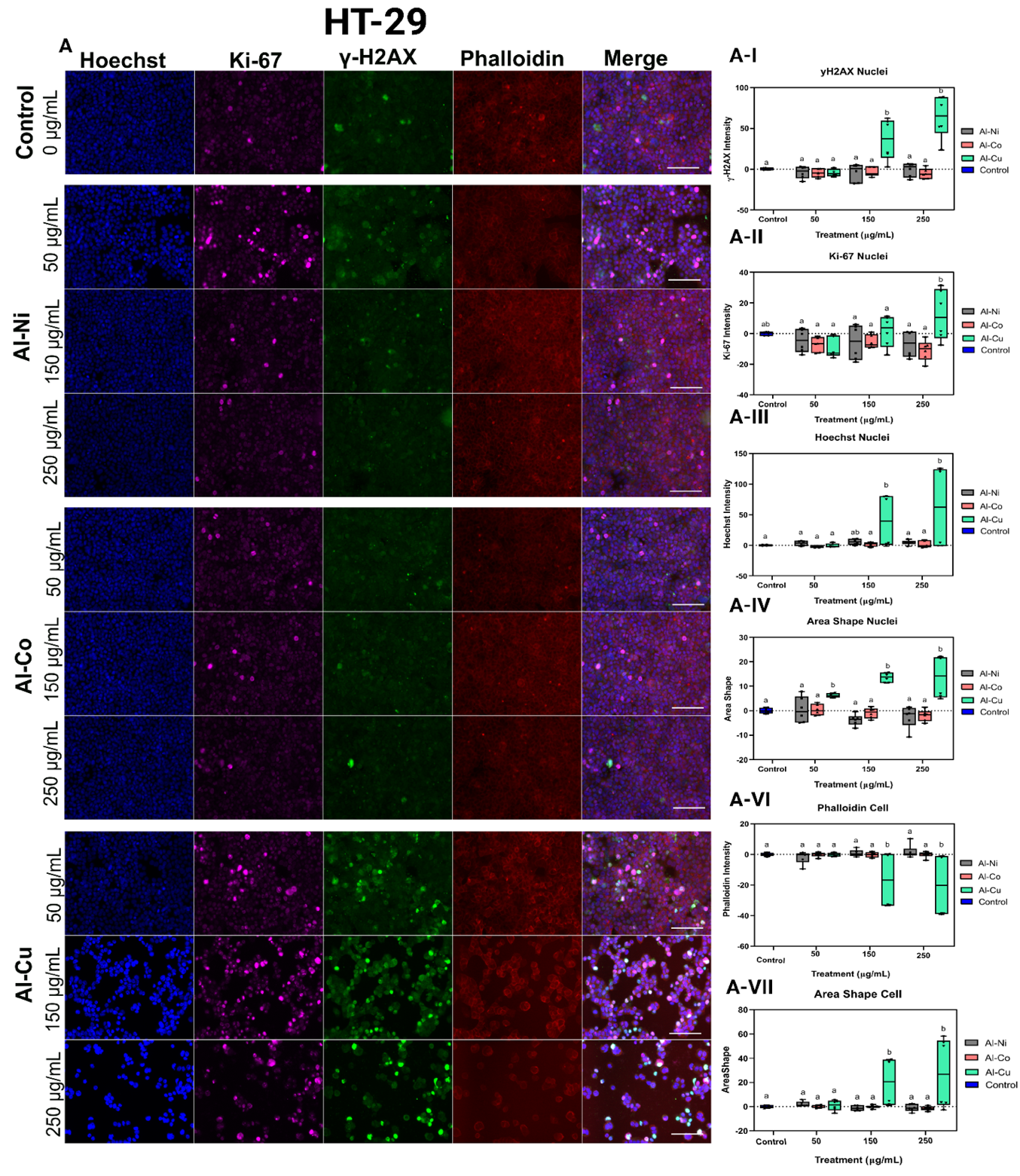

**Figure SI-10. Immunocytochemical and morphometric analysis of HT-29 cells exposed to LDH nanoparticles.** HT-29 cells were treated for 24 h with Al–Ni, Al–Co, and Al–Cu layered double hydroxide (LDH) nanoparticles at 50, 150, and 250 µg/mL. Cells were stained with Hoechst 33342 (blue, nuclei), Ki-67 (magenta, proliferation-associated marker), γ-H2AX (green, DNA damage-associated marker), and phalloidin (red, actin cytoskeleton). Representative fluorescence images are shown together with the corresponding quantitative analyses of fluorescence intensity and morphometric parameters (A-I to A-VII). Scale bar = 100 µm.
